# Nanoscale organization of ryanodine receptor distribution and phosphorylation pattern determines the dynamics of calcium sparks

**DOI:** 10.1101/2021.10.19.465028

**Authors:** M. Hernández Mesa, J. van den Brink, W.E. Louch, K.J. McCabe, P. Rangamani

**Author notes:** To whom correspondence must be addressed.

## Abstract

Super-resolution imaging techniques have provided a better understanding of the relationship between the nanoscale organization of function of ryanodine receptors (RyRs) in cardiomyocytes. Interestingly recent data have indicated that this relationship is disrupted in heart failure (HF), as RyRs are dispersed into smaller and more numerous clusters. However, RyRs are also hyperphosphorylated in this condition, and this is reported to occur preferentially within the cluster centre. Thus, the combined impact of RyR relocalization and sensitization on Ca^2+^ spark generation in failing cardiomyocytes is likely complex and these observations suggest that both the nanoscale organization of RyRs and the pattern of phosphorylated RyRs within clusters could be critical determinants of Ca^2+^ spark dynamics. To test this hypothesis, we used computational modeling to quantify the relationships between RyR cluster geometry, phosphorylation patterns, and sarcoplasmic reticulum (SR) Ca^2+^ release. We found that RyR cluster disruption results in a decrease in spark fidelity and longer sparks with a lower amplitude. Phosphorylation of some RyRs within the cluster can play a compensatory role, recovering healthy spark dynamics. Interestingly, our model predicts that such compensation is critically dependent on the phosphorylation pattern, as phosphorylation localized within the cluster center resulted in longer Ca^2+^ sparks and higher spark fidelity compared to a uniformly distributed phosphorylation pattern. Our results strongly suggest that both the phosphorylation pattern and nanoscale RyR reorganization are critical determinants of Ca^2+^ dynamics in HF.

**Significance Statement:** RyRs are ion channels located on the membrane of the sarcoplasmic reticulum that are responsible for an increase in cytosolic Ca^2+^ during cell excitation. Here, we investigate how the geometry of RyR clusters combined with spatial phosphorylation patterns impacts on Ca^2+^ spark generation and kinetics. The findings from our study show that both phosphorylation pattern and RyR cluster shape and dispersion have implications on Ca^2+^ spark activity and provide insights into altered Ca^2+^ dynamics during HF.

## 1 Introduction

Excitation-contraction coupling (ECC) refers to a series of electrochemical and mechanical processes that repeat during each heartbeat, and allow coupling between electrical excitation and contraction of the heart (*1*). During electrical depolarization of a cardiomyocyte, voltage-gated L-type Ca^2+^ channels open, leading to an influx of Ca^2+^. This incoming Ca^2+^ triggers additional Ca^2+^ release from the sarcoplasmic reticulum (SR) through ryanodine receptors (RyRs) located on the SR membrane (Figure 1). This process of Ca^2+^-induced Ca^2+^ release is central to ECC, as binding of cytosolic Ca^2+^ to the myofilament protein troponin C triggers contraction. Relaxation then occurs as Ca^2+^ levels decline due to RyR closure, Ca^2+^ recycling into the SR by the SR Ca^2+^- ATPase (SERCA) pump, and Ca^2+^ flux out of the cell through the Na/Ca exchanger and the plasma membrane Ca^2+^-ATPase pump.

**Figure 1:**
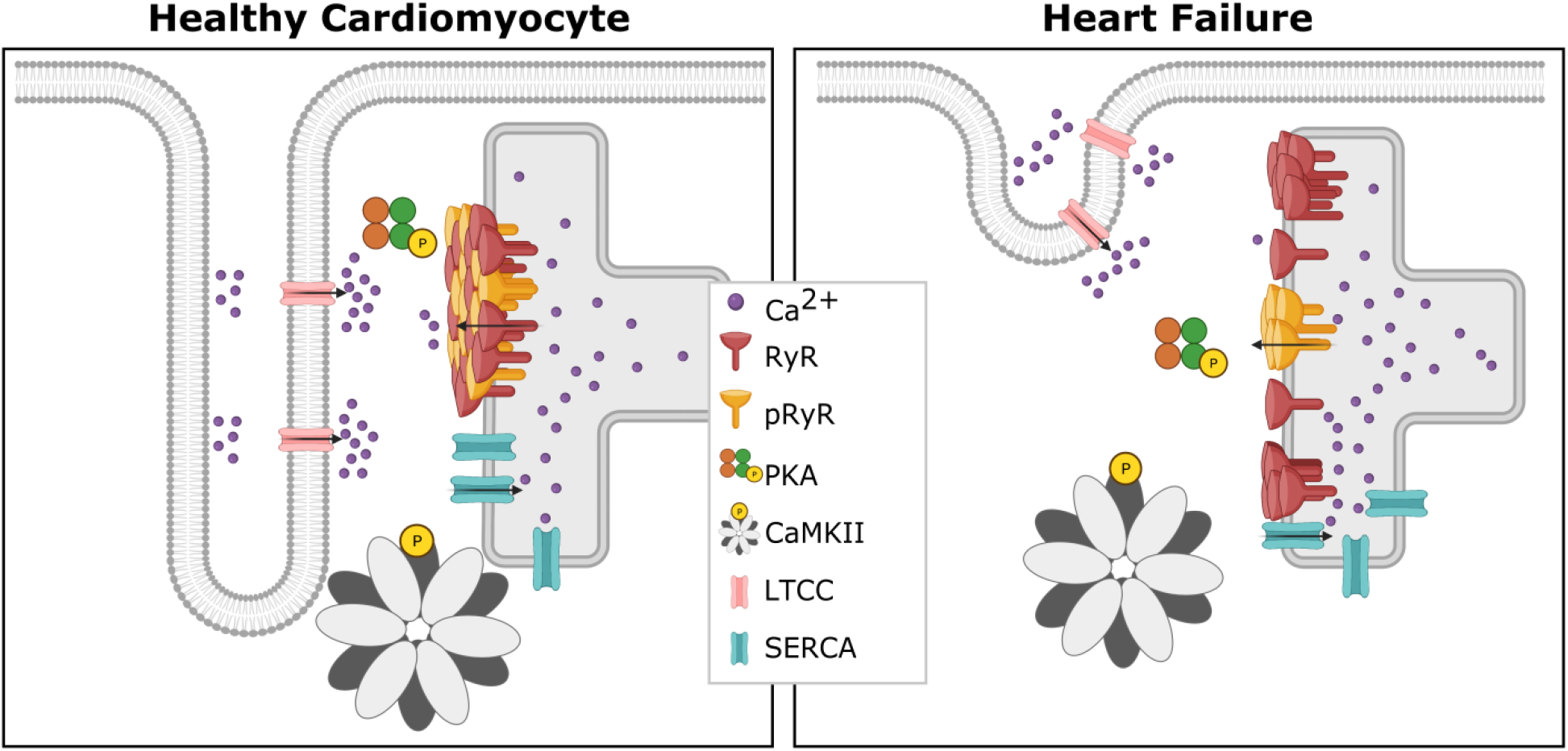
Ca^2+^-induced Ca^2+^ release occurs at units called dyads, where t-tubules and SR are in close proximity. In comparison with healthy cardiomyocytes (left), diseases such as HF have been linked to marked subcellular remodeling (right). Reported changes in failing myocytes include loss of T-tubule density, dispersion of RyR clusters, and changes in the spatial pattern of RyR phosphorylation.

Given the central role of RyRs in controlling cardiomyocyte Ca^2+^ homeostasis and contraction, it is essential that this channel’s function is carefully elucidated. It is known that multiple RyRs collaborate to generate Ca^2+^ sparks, which are the fundamental units of SR Ca^2+^ release. While the precise number of RyRs that produce a spark has been debated, it is generally accepted that 6-20 channels participate, yielding a release event 10-15 ms in duration (*2*). This cooperative activity appears to be enabled by the organization of RyRs into clusters on the SR membrane, which allows for cooperative activation. Nearby clusters of RyRs may also collaboratively generate sparks if they are close enough together (<150 nm) to enable rapid Ca^2+^ diffusion between them (*3–5*). Such functional groupings of neighbouring RyR clusters are commonly referred to as Ca^2+^ release units (CRUs) (*5, 6*).

HF is often characterized on the cellular level by a marked loss of t-tubules and the creation of “orphaned” RyRs. However, recent data from super-resolution imaging studies have revealed that the morphology of CRUs also changes in this condition, resulting in smaller and more numerous RyR clusters (*4, 7–9*) (Figure 1). This RyR cluster dispersion has been linked to slower Ca^2+^ spark kinetics and a resulting desynchronization of the overall Ca^2+^ transient (*7, 10*). However, RyR activity is also critically regulated by phosphorylation by protein kinase A (PKA), Ca^2+^ calmodulin kinase type II (CaMKII), and various phosphatases (*11*) (Figure 1). It is well known that phosphorylation increases RyR open probability (*11*), and that RyR phosphorylation is augmented during HF (*2, 12, 13*). Therefore, understanding RyR function and dysfunction during HF requires an integrated understanding of nanoscale RyR localization and phosphorylation status. Complicating the issue is the finding that RyR phosphorylation patterns may not be uniform. Indeed, Sheard *et al.* (*8*) reported while RyRs are uniformly phosphorylated across clusters in healthy cardiomyocytes, cells from failing hearts exhibited a higher density of PKA-phosphorylated RyRs at the center of RyR clusters. The functional implications of these changing phosphorylation patterns are unclear, and have not been addressed in previous computational studies which assumed that all RyRs in a cluster are equally phosphorylated (*14–16*).

In the present work, we investigated how the spatial organization of RyR clusters affects Ca^2+^ dynamics, with a particular focus on changing patterns of phosphorylation. To this end, we have adapted an existing mathematical model of the CRU (*7*) to include distinct Ca^2+^ sensitivities of individual RyRs. We expect that our predictions will motivate the design of experiments that can decipher how these localized, nanoscale relationships contribute to impaired Ca^2+^ homeostasis during HF.

## 2 Methods

### 2.1 Model development

The mathematical model from the work of Kolstad *et al.* (*7*) was adapted to differentiate between two subgroups of RyRs: phosphorylated and non-phosphorylated RyRs. As described in (*7*), the model was extended from the model of Hake *et al.* (*17*) and includes a stochastic model of RyR opening developed by Cannell *et al.* (*18*) for both phosphorylated and unphosphorylated populations. The system of partial differential equations applied for the spatio-temporal evolution of [Ca^2+^] in the cytosolic domain Ω_*c*_ was:

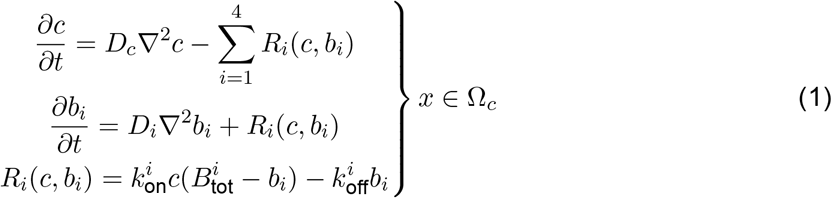

with *c* corresponding to the calcium concentration in the cytosolic domain, *D*_*c*_ the diffusion constant of calcium, *b*_*i*_ the concentration of the corresponding buffer and 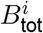 the corresponding total buffer concentration. In the cytosolic domain the buffers ATP, calmodulin, troponin and Fluo-4 were included and correspond respectively to numbers 1 to 4 in the equations listed in Equation 2. For the SR domain Ω_*s*_ (both junctional and non-junctional SR) one calsequestrin buffer was included and its concentration will be denoted by *b*_5_. The following equations correspond to the spatio-temporal evolution of [Ca^2+^] in the SR domain *Ω*_*s*_:

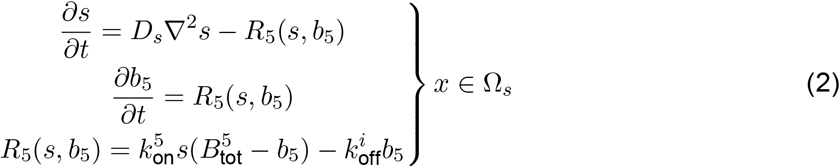

with *s* corresponding to the calcium concentration in the SR domain. Both domains were coupled at the SR membrane with the following boundary conditions:

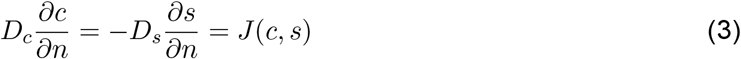

with

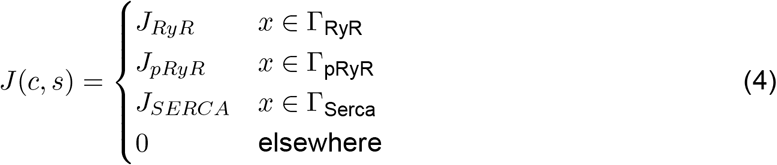

The SERCA flux formulation was taken from the three-state SERCA model from Tran *et al.* (*19*). The *J*_*RyR*_ and *J*_*pRyR*_ formulations were identical to the *J*_*RyR*_ flux formulation from Kolstad *et al.* (*7*), where a two state stochastic model was used:

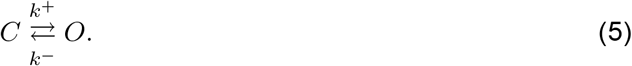

Here, C describes the conductive state and O the non-conductive state. The *k*^−^ and *k*^+^ variables were defined as:

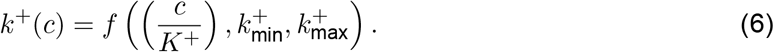

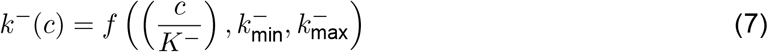

The difference between *J*_*RyR*_ and *J*_*pRyR*_ is the value used for *K*^+^. Since RyR phosphorylation sensitizes the channel more to Ca^2+^ (*12*), we simulate this by lowering the *K*^+^ value. For no phosphorylation conditions the *K*^+^ value was 55 μM. For phosphorylated RyRs the *K*^+^ value was 25 μM, if a non homogeneous phosphorylation pattern was considered and 45 or 35 μM if a blanket phosphorylation was assumed. Supplementary Tables 1 and 2 show the model and buffering parameters.

### 2.2 Geometries

The model consists of a single CRU containing both cytosolic and SR domains (Ω_*c*_ ∩ Ω_*s*_). We assumed that the simulated CRU was a 3D cube with volume 1.008 μm × 1.008 μm × 1.008 μm. Cube dimensions were 12 nm × 12 nm × 12 nm (Δ*x* = 12 nm) with 84 voxels per axis.

The RyR geometries were inspired by the super-resolution images of Kolstad *et al.* (*7*). However, in this work we constrained the number of RyRs to 50 for all geometries, which is within the measured range in experiments (*7*). Five different geometries were designed. The first two geometries (G1 and G2) contain a single cluster to simulate the healthy case (Figure 2A). The main difference between G1 and G2 is the shape of the cluster; while G1 is a more compact distribution, G2 is oblong. By comparing G1 and G2, we can determine whether altering cluster shape without adjusting the density has an impact on spark dynamics. The latter 3 geometries (G3, G4, and G5) differ in the number of CRU sub-clusters. Disrupted RyR clusters containing several sub-clusters have been observed in rats with HF (*7*). Therefore G3, G4, and G5 contain 50 RyRs distributed in 2, 3, and 12 sub-clusters, respectively.

**Figure 2:**
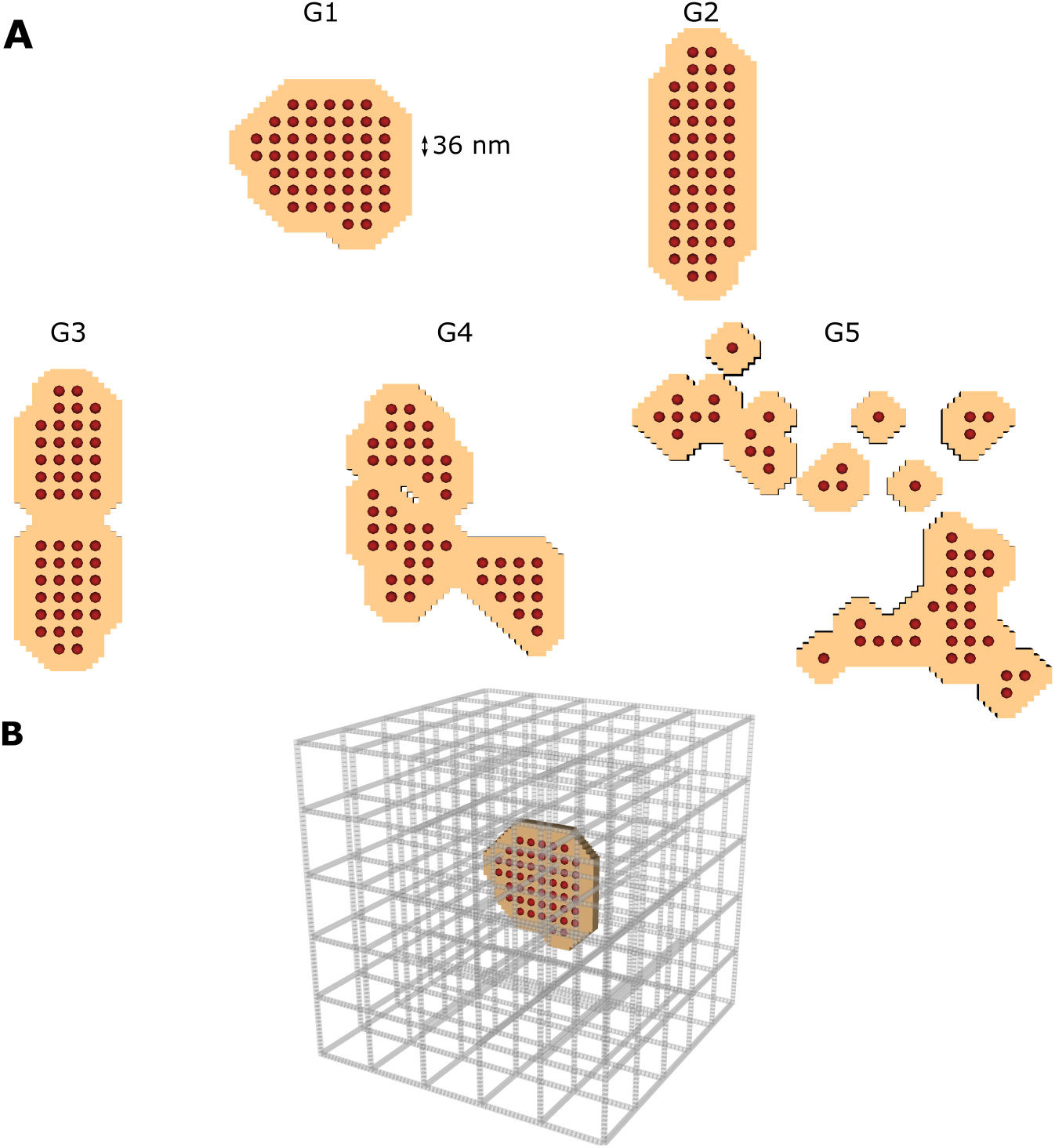
(A) The five different RyR geometries studied in the model. The red dots show the single RyRs and the tan coloured area represents the junctional SR membrane. All five geometries contain 50 RyRs. The first two geometries (G1 and G2) contain a single cluster to simulate the healthy case. The latter 3 geometries (G3, G4, and G5) differ in the number of CRU sub-clusters, to simulate the disrupted RyR clusters observed during HF. G3, G4 and G5 are organized into 2, 3 and 12 sub-clusters respectively. (B) The computational domain of the cytosol is presented. The space corresponds 1.008 μm × 1.008 μm × 1.008 μm. The grey bars represent the non-junctional SR.

Each RyR has an area of 36 nm×36 nm. The center-to-center distance between two neighbouring RyRs is 36 nm, as described in (*7*). The jSR was designated as the area of a single receptor around each RyR (’padding’), in the same fashion as Kolstad *et al.* (*7*). The SR surface area is given in Supplemental Table 3. The nSR was assumed to be a regular grid throughout the cytosol. The SERCA surface has a range of 4.54 4.85 μm^2^. The jSR and nSR surface areas agree with the surface areas measured by Hake *et al.* (*17*) in EM tomography dyad reconstructions. Figure 2B shows the location of the RyRs and jSR within the nSR grid for the G1 geometry.

### 2.3 Phosphorylation patterns

Since the goal of this study is to analyze the impact of spatial phosphorylation patterns on Ca^2+^ spark signals, three different phosphorylation patterns were applied (Figure 3). Additionally, simulations with no phosphorylated RyRs and with blanket phosphorylation of all RyRs (to emulate previous computational models) were carried out. This resulted in 8 phosphorylation setups per geometry. An example of the 8 phosphorylation setups for geometry G1 is shown in Figure 3.

**Figure 3:**
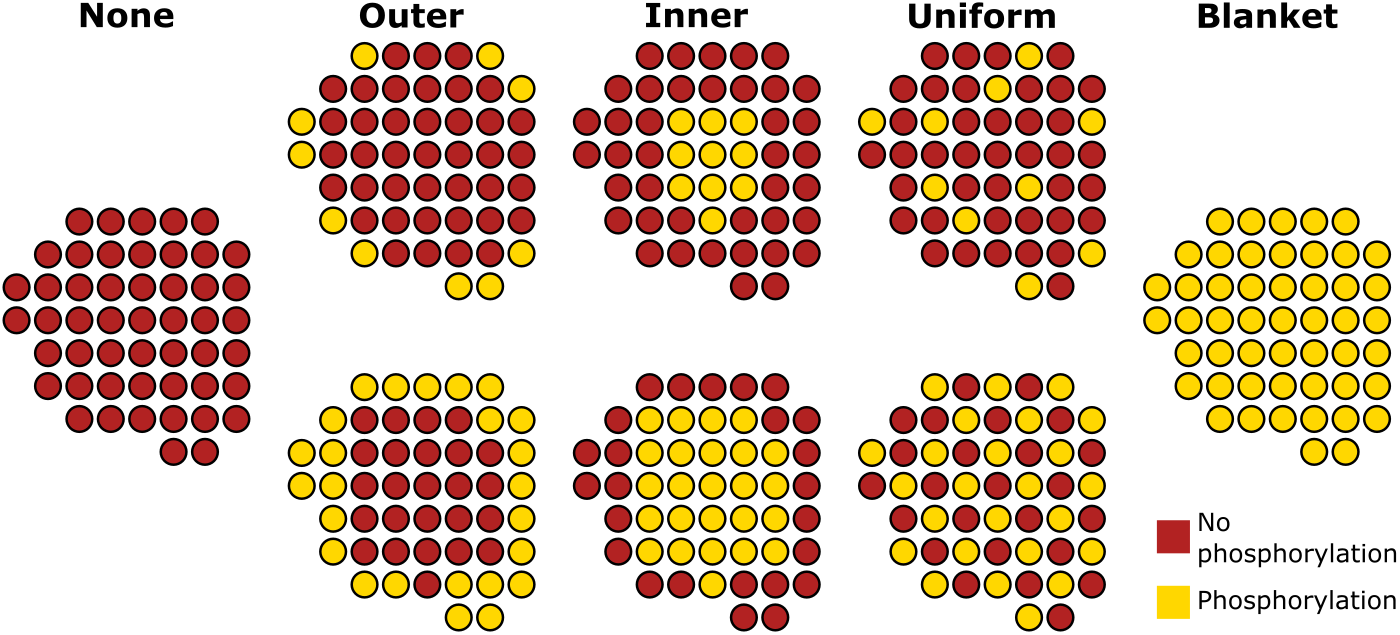
Schematic of the different phosphorylation setups used in the model demonstrated using the geometry G1 as an example case. For the three phosphorylation patterns (outer, inner and uniform) two configurations were assumed: 20% (upper row) and 50% (lower row) of the RyRs being phosphorylated.

Sheard et al. (*8*) reported that in HF, a spatial gradient of RyR hyperphosphorylation occurs with the highest phosphorylation levels occurring within the center of the nanodomain. In order to simulate this, phosphorylated RyRs were chosen using principal component analysis (PCA) (*20*). The eigenvectors of the covariance matrix of the RyR positions were calculated to estimate the directions of maximal information of the CRU. With both orthogonal eigenvectors, an ellipse is generated. The ellipse dimensions were then decreased until the desired number of RyRs was phosphorylated. Figure S1 depicts an example of the PCA method applied to geometry G1 to calculate the inner phosphorylation. By applying this technique we select the desired number of RyRs in the center of the CRU when simulating HF conditions. We also chose to study the opposite pattern using the same PCA technique but phosphorylating RyRs near the outer boundary of the cluster. Although this specific phosphorylation pattern has not been observed in experimental studies, we include it in our simulations for completeness. These two PCA-based phosphorylation schemes will be referred to as inner and outer phosphorylation patterns from here onwards. For each pattern, two phosphorylation levels were simulated: 20% (10 RyRs) and 50% (25 RyRs) phosphorylation of the cluster. Finally, in order to simulate a uniform distribution of the phosphorylation pattern, as may be the case in the healthy condition (described in (*8*)), we simulated a uniformly distributed phosphorylation condition (again at 20% and 50% levels). In total, 8 different phosphorylation configurations per geometry were analyzed, all of which are visualized for the G1 geometry in Figure 3. An overview of the phosphorylation patterns used for G2, G3, G4 and G5 is provided in the supplementary material (Figures S2 and S3). Since the 8 different phosphorylation patterns were simulated for 5 different geometries, this results in 40 different spatial configurations.

### 2.4 Numerics

Since the model used assumes a stochastic model for RyR opening, 200 runs were carried out for each of the 40 configuration setups (5 geometries × 8 phosphorylation patterns). The simulations were initiated by randomly opening 1 RyR within the specific geometry, and the RyR fluxes were then calculated. Opening and closing of a single RyR was based on local Ca^2+^ concentrations. The simulations were terminated when all RyRs were closed and remained closed for 1 ms. Therefore, we define spark duration as the duration of the simulations 1 ms after all RyRs are closed. The stochastic RyR model was calculated for a fixed time step of Δ*t* = 1 ms. The model calculations were very stiff due to the small element volumes and the large fluxes. Therefore, the calculations of the fluxes in the model were solved analytically as described in (*7*). We also used the following definitions to classify and quantify our results. We consider a spark to be successful if local increases in simulated Ca^2+^ fluorescence intensity exceeded 30% (Δ*F* /*F*_0_ ≥ 0.3), and used the fraction of these successful sparks to estimate spark fidelity. The confidence interval of spark fidelity was calculated using the Agresti-Coull confidence interval (*21*). Additional spark properties including the amplitude, time to peak (TTP), and spark simulation duration were used to compare and contrast different scenarios. For these three properties, the confidence interval was calculated using the standard error.

## 3 Results

### 3.1 Circular CRUs increase spark fidelity and amplitude

First, simulations without RyR phosphorylation were carried out for all 5 geometries. Figure S4 shows that our simulation results match the results from Kolstad *et al.* (*7*), as spark fidelity and amplitude decrease with increasingly dispersed RyR configurations. We additionally compared two single-cluster geometries (G1 and G2), where G1 had a compact circular geometry and G2 a more oblong arrangement. Interestingly, a clear difference in spark fidelity (ie. proportion of sparks with ΔF/F_0_ > 0.3) is observed between these two geometries (Figure S4). This is illustrated by the spark time courses for G1 *versus* G2 (Figure 4A and B, respectively). For G1, 63 out of 200 sparks were successful (dark gray sparks), while for G2 only 29 successful sparks occurred. Figure 4C shows the probability of spark generation for both geometries. Out of 200 simulations, 31.8% *±* 6.4% surpassed the spark detection threshold for G1. This number was significantly lower in the G2 geometry, which had a spark fidelity of 15.2% ± 4.9%. In addition to a decrease in fidelity, the oblong geometry in G2 also results in a significantly lower amplitude (Figure 4D) and longer time to peak (TTP), see Figure 4E. A scatter plot of these measurements in individual sparks is shown in Figure 4F.

**Figure 4:**
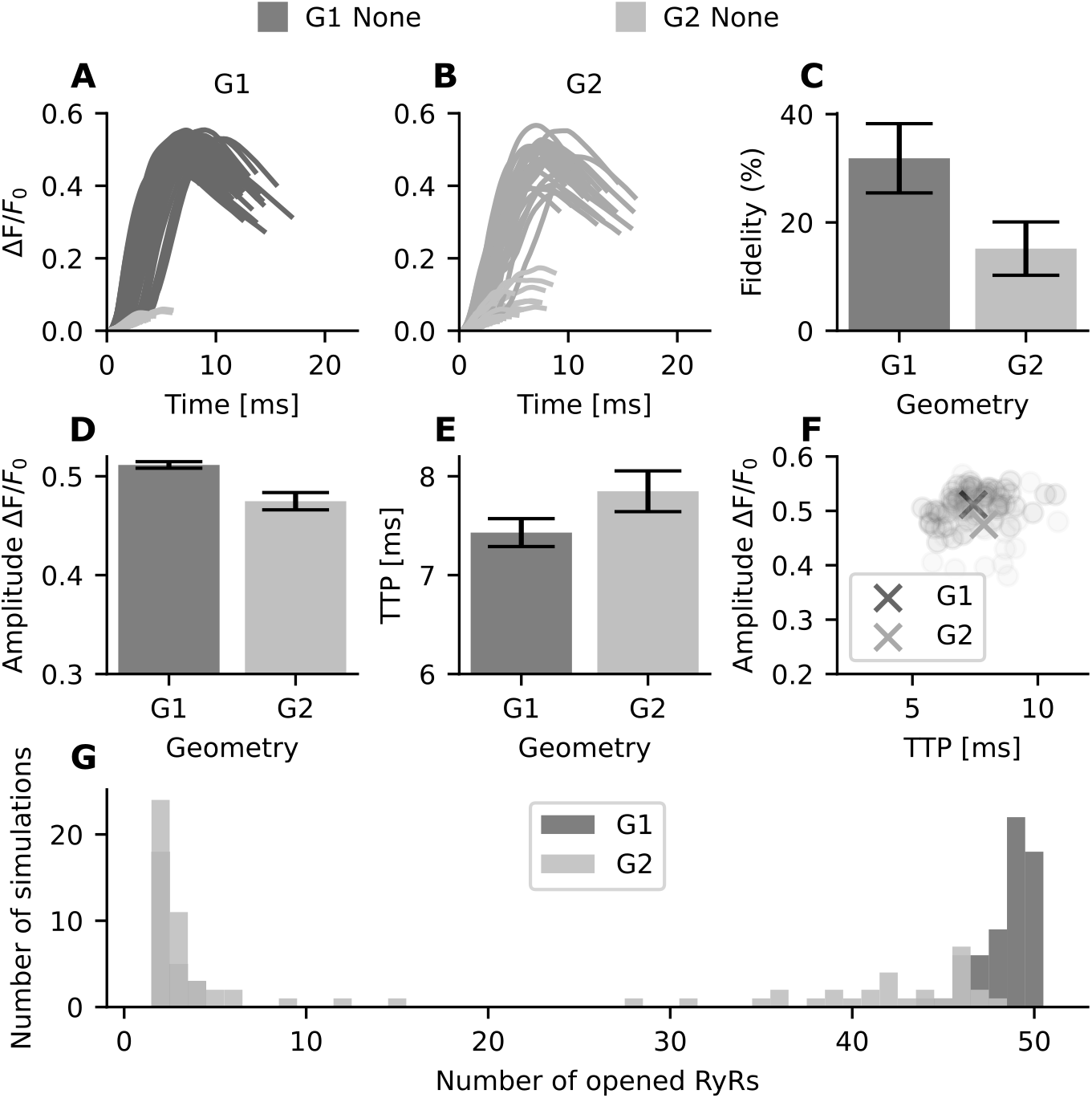
Spark properties of unphosphorylated RyR in geometries G1 and G2. For each geometry 200 simulations were conducted. (A) Intensity over time for G1 calcium sparks. dark gray: successful sparks (n=63), light gray: failed sparks. (B) Intensity over time for G2 calcium sparks. dark gray: successful sparks (n=29), light gray: failed sparks. (C) The probability of spark generation (spark fidelity) for G1 and G2. Error bars indicate the 95% Agresti-Coull confidence interval. (D) Spark amplitude (mean ± standard error) for successful sparks in G1 and G2. (E) Average time to peak (TTP) (mean ± standard error) for G1 and G2, in ms. (F) Scatter plot of spark amplitude *versus* TTP. The crosses represent the mean values across all simulations for G1 and G2. (G) Histogram tracking total number of opened RyRs across individual simulations for G1 and G2. For example, for geometry G1 all 50 RyRs are opened in 18 simulations, whereas for G2 no simulations had 50 open RyRs.

We next investigated, whether the lower amplitude, slower kinetics, and lower fidelity of Ca^2+^ sparks generated by the oblong G2 vs compact G1 configuration were linked to a differing number of RyRs being activated. To this end, we plotted a histogram of the number of RyRs opened per simulation (Figure 4G). Note that the G1 geometry exhibits an “all-or-none” behaviour, with bimodal clustering near either one or all 50 of the RyRs being activated in a single simulation. In comparison, G2 shows more varied behavior, and no simulations with full activation. This behaviour is also observed in the activation map of a single simulation for both G1 and G2 (Figure S5). For the G2 simulation activation map shown, the RyRs located at the edges of the geometry are not activated. Thus, our results show that the nanoscale organization of the unphosphorylated RyR clusters affects Ca^2+^ dynamics, as compact, circular nanoclusters generate larger and faster Ca^2+^ sparks than elongated nanoclusters.

### 3.2 Phosphorylation pattern of the RyR cluster determines spark properties

As shown in (*8*), different spatial patterns can be observed for RyR phosphorylation in the CRU. However, computational models generally assume all RyRs in a CRU to be identically sensitized due to a lack of spatial detail (*14–16*). We next tested if there was a difference in Ca^2+^ dynamics when all receptors are phosphorylated (”blanket” phosphorylation) as opposed to some percentage of uniformly phosphorylated receptors in the cluster (see Figure 3 for schematic). To capture the effects of receptor phosphorylation and calcium sensitivity, we varied *K*^+^ in the range [25 μM, 35 μM, 45 μM, and 55 μM]. Here 55 μM represents nonphosphorylated RyR and therefore 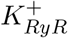. On the other hand, 25 μM represents the situation of uniform phosphorylation (both 20% and 50%); 35 μM and 45 μM represent blanket phosphorylation scenarios. These three values represent 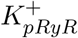.

Sparks generated by a 20% uniform phosphorylation pattern (with 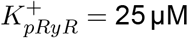 and 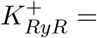 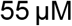) were compared to blanket phosphorylation, with 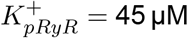. A pattern of 50% uniform phosphorylation (with 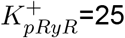 and 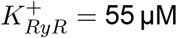) was in turn compared with blanket phosphorylation 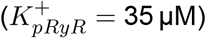. The results are shown for G2 and G3 in Figure 5. The lighter shades represent a 20% uniform phosphorylation pattern (light red) and blanket phosphorylation with a *K*^+^ value of 45 (light gray). The darker shades represent a 50% uniform phosphorylation pattern (dark red) and blanket phosphorylation with a *K*^+^ value of 35 (dark gray). We find that, by adjusting *K*^+^ values, outputs from blanket phosphorylation match those from uniform phosphorylation for all analyzed spark properties (Figure 5A-D).

**Figure 5:**
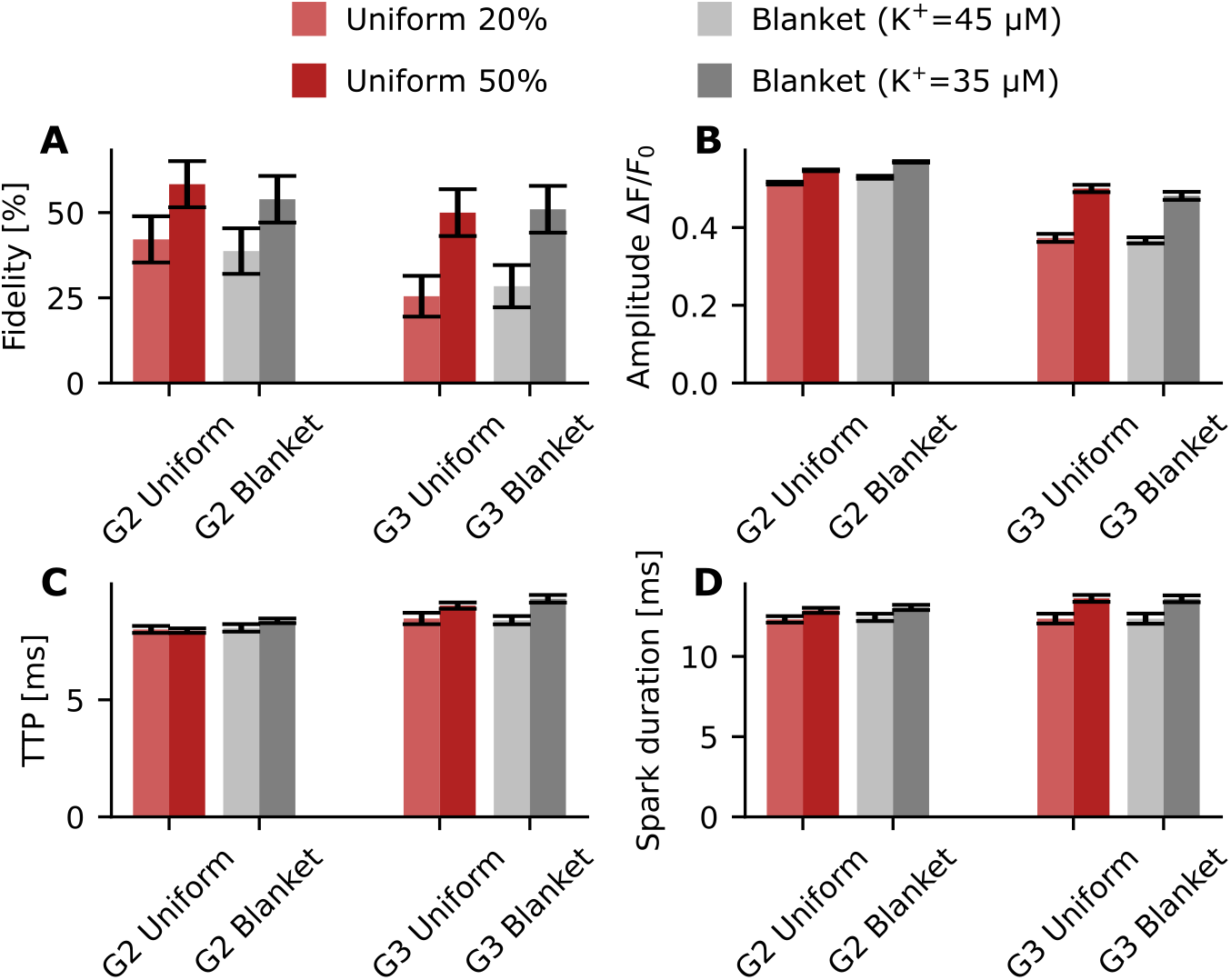
Properties of sparks obtained from uniformly distributed phosphorylation pattern versus blanket phosphorylation for G2 and G3. Red bars represent simulations with a uniformly distributed phosphorylation pattern of 20% (light red) or 50% (dark red) of the receptors. The gray bars represent simulations with a blanket phosphorylation pattern. (A) Fidelity for G2 and G3 at different phosphorylation levels and patterns; the black bars indicate the 95% Agresti-Coull confidence interval. (B) Amplitude (mean *±* standard error) is shown using a bar chart (for G2 uniform 20% n=84 (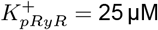 and 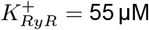), for G2 uniform 50% n=117 (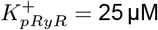 and 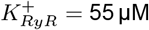), for G2 blanket 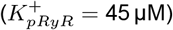 n=77, for G2 blanket 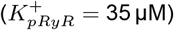 n=108, for G3 uniform 20% n=50 (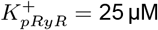 and 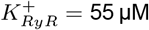), for G3 uniform 50% n=100 (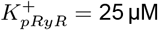 and 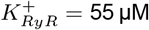), for G3 blanket 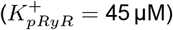 n=56, and for G2 blanket 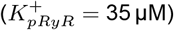 n=102. (C) TTP (mean *±* standard error) is shown using a bar chart. (D) Spark duration (mean *±* standard error) is shown using a bar chart.

We next evaluated the effect of different phosphorylation patterns of the RyR nanocluster in G1 (Figure 6). RyR phosphorylation leads to an increase in Ca^2+^ spark fidelity regardless of pattern (Figure 6A). When comparing the different patterns at the same degree of phosphorylation (20% or 50%), inner phosphorylation results in the largest increase in fidelity, followed by the uniform pattern, with the outer pattern exhibiting the smallest increase (62.3% ± 6.7% vs 60.3% ± 6.7% vs 58.8% ± 6.8% at 50% phosphorylation , and 48.5% ± 6.9% vs 47.1% ± 6.9% vs 31.8% ± 6.4% at 20% phosphorylation).

**Figure 6:**
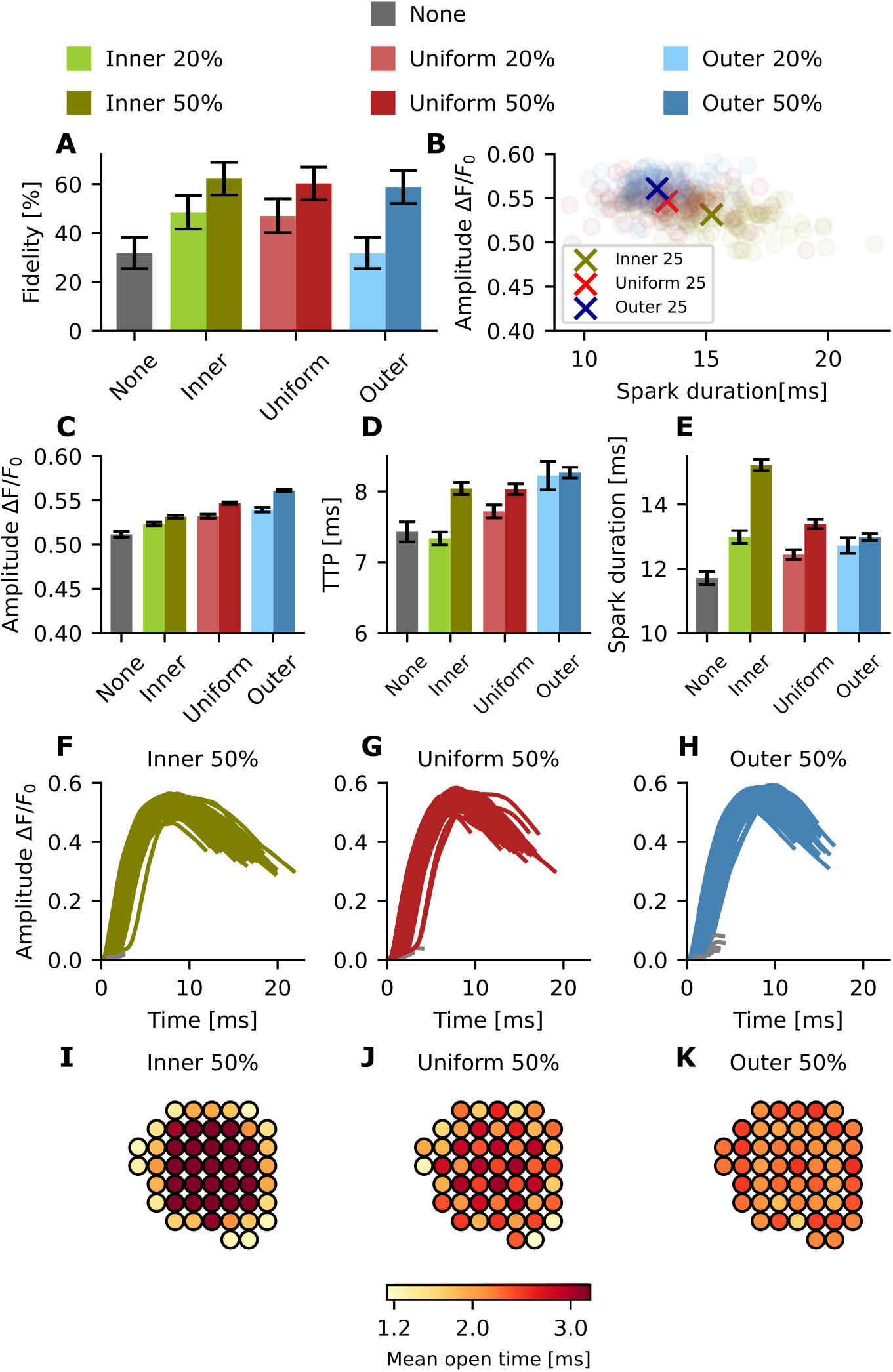
Effect of phosphorylation pattern on Ca^2+^ spark properties in G1. For each pattern, 200 simulations were conducted. green: inner, red: uniform, blue: outer. Lighter shades: 20% phosphorylation, darker shades: 50% phosphorylation. (A) Spark fidelity for inner, uniform, and outer phosphorylation patterns in G1. The black bars indicate the 95% Agresti-Coull confidence interval. (B) Scatter plot comparing spark duration and amplitude. Circles: single simulations, crosses: mean values. (C) Spark amplitude (mean ± standard error) for inner, uniform, and outer phosphorylation patterns in G1. (For no phosphorylation n=63, for inner 20% n=97, for inner 50% n=125, for uniform 20% n=94, for uniform 50%=121, for outer 20% n=63, and for outer 50% n=118) (D) TTP (mean ± standard error) for inner, uniform, and outer phosphorylation patterns in G1. (E) Spark duration (mean ± standard error) for inner, uniform, and outer phosphorylation patterns in G1. (F) Intensity timecourse for successful sparks with inner phosphorylation. (G) Intensity timecourse for successful sparks with uniform phosphorylation. (H) Intensity timecourse for successful sparks with outer phosphorylation. (I) Mean open time for each RyR throughout 200 simulations - inner 50% phosphorylation. (J) Mean open time for each RyR throughout 200 simulations - uniform 50% phosphorylation. (K) Mean open time for each RyR throughout 200 simulations - outer 50% phosphorylation.

Plotting spark duration against spark amplitude (Figure 6B), we found that outer phosphorylation produces slightly larger sparks than uniform or inner phosphorylation (ΔF/F_0_ =0.560 ± 0.001 for 50% outer phosphorylation vs 0.547 ± 0.001 for 50% uniform phosphorylation vs 0.531 ± 0.002 for 50% inner phosphorylation, Figure 6C). We also measured TTP and spark duration for the same phosphorylation patterns (Figures 6D and 6E). Phosphorylation leads to increased spark duration in our model, across all patterns. An increased TTP was also seen across the different patterns, with the exception of the inner 20% configuration. For 20% phosphorylation, outer phosphorylation leads to the highest increase in TTP, followed by uniform, then inner (Figure 6D). However, for 50% phosphorylation there is no significant difference between the TTP for the inner and uniform patterns (around 8 ms). This suggests that the relative effect on TTP decreases with increasing fraction of phosphorylated RyRs.

To mechanistically understand the rather complicated dependence of spark kinetics on phosphorylation pattern, we studied the dynamics of individual simulations by tracking the opening and closing of the RyRs for single representative sparks (see movies M1-M6). Based on these movies, we first confirm that with inner phosphorylation, the TTP is faster compared to other phosphorylation patterns (Movie M4). However, due to the phosphorylation, the inner RyRs are more likely to reopen, which prolongs the total spark duration and creates a plateau of Ca^2+^ release, an effect that is particularly prominent at high phosphorylation levels. Additionally, the sensitized RyRs are able to sustain the regenerative release longer, resulting in higher fractional release from the SR. Therefore, the morphology of the spark changes from containing a single discernable peak in the unphosphorylated case to a plateau phase in the phosphorylated cases. The sparks measured with an inner phosphorylation pattern have a longer duration compared to the uniform and outer case (15.2 ± 0.18 ms vs 13.4 ± 0.14 ms vs 13.0 ± 0.11 ms at 50% phosphorylation, and 13.0 ± 0.19 ms vs 12.4 ± 0.15 ms vs 12.7 ± 0.24 ms at 20% phosphorylation), despite the shorter TTP (see Figure 6F-H). Importantly, phosphorylation patterns were found to have a large impact on the mean open time of individual RyRs within the cluster (Figure 6I-K). With a uniform phosphorylation pattern, RyRs located near the center of the cluster tended to be open longer than the channels located at the outer boundary (Figure 6J). In other words, the central channels anchored regenerative release. This effect is emphasized even more strongly with an inner phosphorylation pattern, since the inner channels are sensitized both by their spatial location and their phosphorylated state (Figure 6I). For the outer phosphorylation pattern, the effect is reversed, with channels across the cluster exhibiting close to uniform mean open times (Figure 6K). Taken together, our simulations predict that the phosphorylation pattern within the CRU has a marked impact on Ca^2+^ spark dynamics, as inner phosphorylation leads on average to higher fidelity, lower amplitudes, and longer spark durations than uniform or outer phosphorylation patterns.

### 3.3 Inner phosphorylation maximizes spark fidelity and increases spark amplitude in a dispersed RyR cluster organization

We next analyzed the effects of RyR phosphorylation in disrupted CRU geometries, which are reported in HF (*7*). As noted above, when all the RyRs are unphosphorylated, we observed a step-wise decrease in spark fidelity and amplitude with increasing dispersal of the CRU geometries (Figure S4). These results match the outcomes of previous work based on the same model (*7*). As described by Sheard *et al.* (*8*), during HF, a shift of the PKA-mediated phosphorylation pattern is observed towards the center of the cluster. Therefore we investigated whether the effects of the phosphorylation patterns shown in Figure 6 are also observed in disrupted geometries.

We first focus on the differences between G2 and G3, since the only difference between these geometries is that G2 contains one cluster and G3 two sub-clusters, while overall cluster shape is maintained (Figure 2). For both geometries, higher spark fidelity is observed when applying inner phosphorylation compared to a uniform or outer phosphorylation (Figure 7A). We also observe a significant increase in spark duration for both geometries with inner phosphorylation in comparison with uniform and outer phosphorylation patterns (14.2 *±* 0.16 ms vs 12.9 *±* 0.16 ms vs 12.76 ± 0.17 ms at major phosphorylation for G2, and 14.7 ± 0.24 ms vs 13.6 ± 0.22 ms vs 13.5 ± 0.24 ms at major phosphorylation for G3) (Figure 7B).

**Figure 7:**
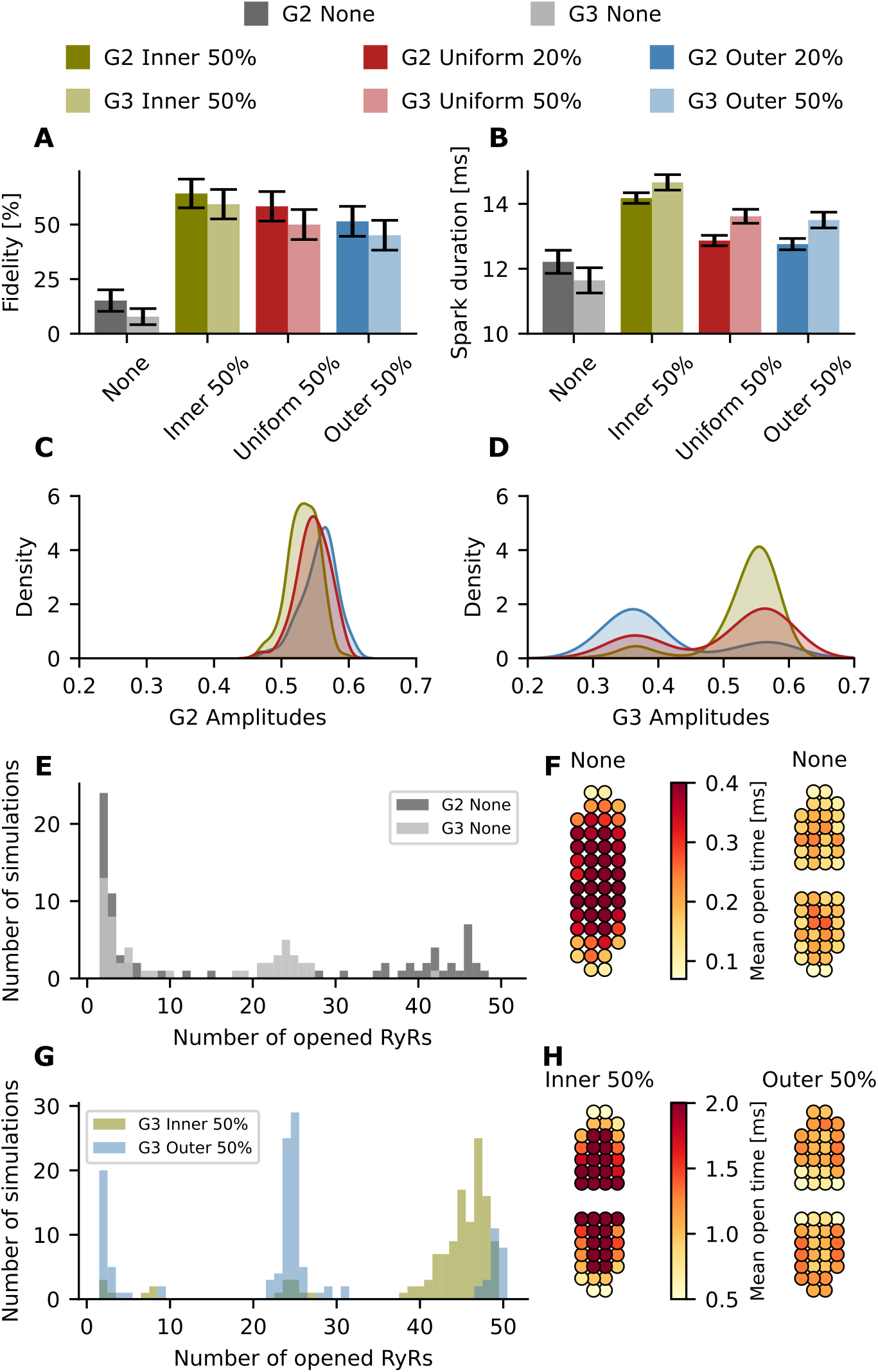
Effect of dispersed and phosphorylated RyR nanoclusters on spark properties. Green: inner; red: uniform; blue: outer; gray:no phosphorylation. Opaque colours: G2; translucent colours: G3. (A) Spark fidelity for none, inner, uniform, and outer phosphorylation in G2 and G3. The black bars indicate the 95% Agresti-Coull confidence interval. (B) Spark duration (mean ± standard error)for none, inner, uniform, and outer phosphorylation in G2 and G3. (C) Kernel density estimate plot of the amplitudes for the three phosphorylation patterns in G2. (D) Kernel density estimate plot of the amplitudes for the three phosphorylation patterns in G3. (E) Histogram of opened RyRs per simulation for G2 and G3 with no phosphorylation. (F) Geometric visualization of mean open times for each RyR throughout all 200 simulations for G2 and G3. (G) Histogram of opened RyRs per simulation for G3 with inner and outer phosphorylation. (H) Geometric visualization of mean open times for G3 with inner and outer phosphorylation.

However, differing effects on spark amplitude were observed for compact geometry G2 and discontinuous geometry G3 (Figure 7C and Figure 7D). Indeed, for G1 and G2, an increase in amplitude is observed when shifting phosphorylation from the inner to the outer pattern (Figures 6 and 7). G3 exhibits the opposite behaviour, as the mean amplitude is highest when assuming inner phosphorylation on amplitude, followed by a uniform phosphorylation and an outer phosphorylation (0.532 ± 0.005 vs 0.500 ± 0.009 vs 0.412 ± 0.010 at major phosphorylation for G3). Interestingly, kernel density plots show that changing the phosphorylation pattern had rather complex effects on spark amplitude, as data from the G2 configuration showed a Gaussian relationship, while biphasic curves were observed for the G3 configuration. To understand the described effects of phosphorylation, histograms showing the number of opened RyRs vs number of simulations (Figure 7E and Figure 7G) are illustrative. When plotting the no phosphorylation case for G2 and G3 in Figure 7E, we observe that in a successful spark for G3, only around 25 RyRs open, as activation was limited to one sub-cluster. For G2, however, we see that if a spark is generated the number of opened RyRs is much higher. It is notable that RyRs in the center of each subcluster tend to open more frequently and for longer durations than RyRs on the boundary (see Figure 7F) resulting in higher probability for the released Ca^2+^ to activate the neighbouring subcluster

When applying a phosphorylation pattern, however, the activation map changed considerably(see Figure 7H). For the inner case, the number of open RyRs increased dramatically (Figure 7G) and phosphorylated RyRs remained open for longer (Figure 7H), leading to more reliable activation of the neighbouring subcluster. This jumping of released Ca^2+^ between subclusters was much less frequent with an outer phosphorylation pattern, with a lower number of RyRs being activated (Figure 7G).

The histogram also helps to explain the biphasic curves in the amplitude kernel density estimate plot (Figure 7D). The first peak, at lower amplitude, represents sparks generated when only one sub-cluster is activated. The second peak, at higher amplitudes, represents sparks generated when both sub-clusters are activated. For inner phosphorylation, the mode occurs at a higher amplitude of around 0.55, whereas outer phosphorylation more often reaches an amplitude of 0.35 more characteristic of single subcluster activation.

We next investigated the effect of phosphorylation on clusters with pronounced dispersion (Figure 8). The activation maps for G4 and G5 are shown in Figure 8A. We found that inner phosphorylation yielded higher spark fidelity pattern than for a uniform or outer phosphorylation pattern (Figure 8B); effects that were more marked than observations made in condensed CRUs (compare with Figure 6). For 50% phosphorylation the fidelity is 38.7 ± 6.69% for an inner phosphorylation, whereas for uniform and outer phosphorylation, the fidelity is to 30.4 ± 6.31% and 12.7 ± 4.57%, respectively. Indeed for G5, this effect is even more relevant since only the inner 50% phosphorylation and the uniform 50% phosphorylation patterns generate any sparks over the 0.3 threshold (Figure 8C). Spark properties calculated for G4 showed that, as for G3, the inner phosphorylation yields longer sparks with a higher amplitude (Figures 8D-F). We do not show the outcomes of the spark analysis for geometry G5 since the number of successful sparks is too low to generate meaningful statistics. These outcomes suggest that phosphorylation pattern may be of particular importance in severely disrupted CRU geometries, where reorganisation of phosphorylated RyRs toward cluster centers may provide a compensatory effect and allow sparks to persist.

**Figure 8:**
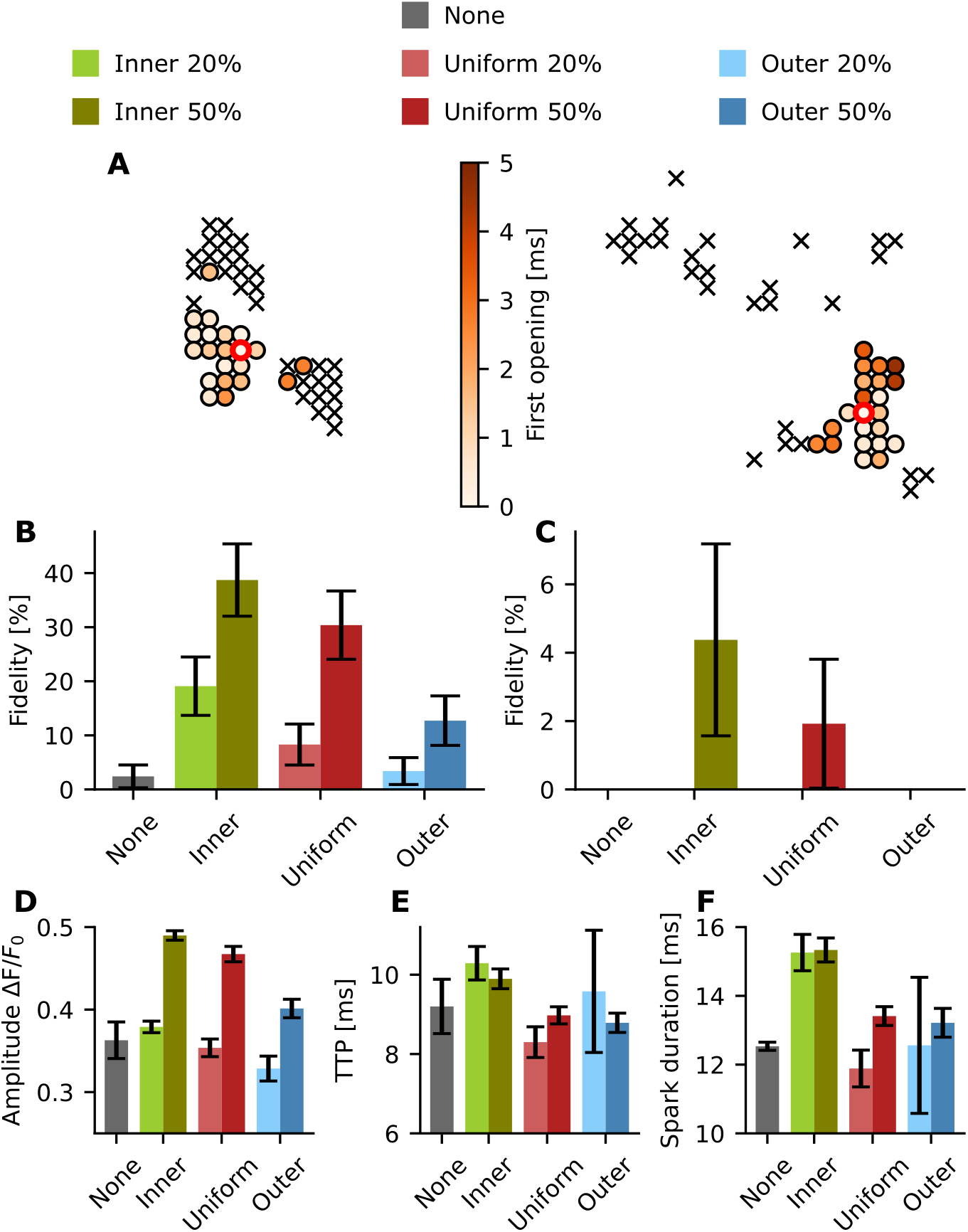
Spark properties for disrupted geometries G4 and G5. For each setup 200 single simulations were conducted. Green: inner; red: uniform; blue: outer; gray:no phosphorylation. light colours: 20% phosphorylation; dark colours: 50% phosphorylation. (A) Activation map with the first opening time of each RyR for a representative simulation for uniform phosphorylation, G4 (left) and G5 (right). ‘x’ indicates that the RyR did not open during the simulation. red circle: initial open RyR. (B) Spark fidelity for none, inner, uniform, and outer phosphorylation in G4. The black bars indicate the 95% Agresti-Coull confidence interval. (C) Spark fidelity for none, inner, uniform, and outer phosphorylation in G5. The black bars indicate the 95% Agresti-Coull confidence interval. (D) Amplitude (mean ± standard error) for none, inner, uniform, and outer phosphorylation in G4 (no phosphorylation n=3, inner 20% n=37, inner 50% n=77, uniform 20%=15, uniform 50%=60, outer 20% n=5, outer 50% n=24). (E) TTP (mean ± standard error) for none, inner, uniform, and outer phosphorylation in G4. (F) Spark duration (mean ± standard error) for none, inner, uniform, and outer phosphorylation in G4.

## 4 Discussion

In this study, we have adapted a stochastic computational model to study the effect of RyR cluster organization and phosphorylation patterns on Ca^2+^ spark dynamics. Using this model, we show that fundamental features of RyR nanoclusters in a CRU, including cluster geometry, phosphorylation pattern, and cluster integrity, interact in combination to regulate spark properties.

As shown in a previous computational study by Walker *et al.* (*22*), spark fidelity in randomly distributed non-square geometries is lower than fidelity in squared geometries. This matches with our observations regarding lower fidelity in less circular geometries, along with lower amplitudes and longer sparks. This can be understood by the fact that in a compact geometry, the RyRs are in closer proximity to other RyRs in the cluster, leading to tighter intra-cluster coupling of all RyRs within the cluster. Indeed, Ca^2+^ leaks out of the sides of the CRU geometry into the cytosol, meaning that RyRs on the outer part of the CRU geometry can experience less Ca^2+^ availability. In oblong clusters, such as the G2 geometry, these effects are amplified at the cluster ends where there are less neighbouring RyRs, leading to reduced coupling. We further observed that more dispersed CRU geometries result in decreased spark fidelity and amplitude, and increased spark duration, consistent with previous computational and experimental work (*7*). These results and the differences between compact and disrupted geometries match the spark property changes measured between healthy and HF patients (*23*).

Based on the redistribution of the spatial pattern observed using enhanced expansion microscopy for the RyR phosphorylation site pSer2808 on Wistar rats undergoing right ventricle HF (*8*), we studied the effect of spatial phosphorylation pattern redistribution on multiple cluster geometries. In our simulations, we observe that inner phosphorylation results in compact distribution of phosphorylated RyRs in the center of the cluster, whereas the outer and uniform patterns can be seen as disrupted phosphorylation subdomains. As seen in this work and in a previous study by Walker *et al.* (*22*), a compact cluster leads to higher fidelity, meaning increased likelihood for RyRs to open and release calcium. This increase in Ca^2+^ makes neighbouring RyRs more likely to open. We find that the average amplitude of the spark is significantly higher when assuming an inner phosphorylation pattern than when assuming a uniform distribution or an outer phosphorylation pattern because there is more consistent activation of adjacent sub clusters. This can be observed in the representative videos included in the supplemental material, where inner phosphorylation leads to an activation of both subclusters, whereas the uniform phosphorylation only activates one of the subclusters. Cluster dispersion leads to low excitability of clusters, and imparied EC coupling fidelity, which is a common property of failing cardiomyocytes. Phosphorylation counteracts this by sensitizing channels, increasing excitability. Our simulations show the reorganization to an inner phosphorylation pattern can further strengthen this compensatory effect.

The phosphorylation of the RyRs is a widely studied and controversial field, since several papers show contradictory results (*24*). Marx *et al.* (*12*) report the dissociation of the FK506 binding proteins (FKBP12.6) from the RyRs after PKA phosphorylation. However, as reported by Bers (*24*), this finding doesn’t match with experiments measured by other groups (*25*). In a different study Marx *et al.* report the impact of FKBP12.6 on coupled gating between neighbouring RyRs (*26*). Based on the results of Marx *et al.*, Sobie *et al.* (*27*) used a computational model to study the impact of coupled gating between RyRs on calcium sparks. Their results justify the decrease in spark fidelity by a disruption of the coupling gating between RyRs. The works of Marx *et al.* and Sobie *et al.* were carried out before super resolution imaging techniques were applied to visualize single molecule localization (e.g. individual RyRs) (*5*). Super resolution imaging has since proven the disruption of RyR clusters during HF (*7*). In a recent study from Asghari *et al.* (*28*), they observe that phosphorylation of the RyRs also leads to disruption of cluster geometries. In their study, the disrupted geometries also show higher spark fidelity and longer full duration half maximum values. These results may seem to contradict the outcomes from Kolstad *et al.* (*7*), where disrupted geometries lead to lower fidelities and lower amplitudes. However, in the study from Asghari *et al.*, healthy compact geometries are compared with phosphorylated disrupted geometries. In our study we also saw that disrupted geometries undergoing phosphorylation could lead to longer and more frequent sparks than compact geometries without phosphorylation (Figure 9). For example, G3 undergoing inner phosphorylation leads to higher fidelities and higher amplitude values than sparks generated from G1 unphosphorylated conditions. Thus, the results from this study are consistent with both the outcomes from Kolstad *et al.* (*7*), when assuming no phosphorylation, and with the results from Asghari *et al.* when comparing compact unphosphorylated geometries with dispersed phosphorylated geometries.

**Figure 9:**
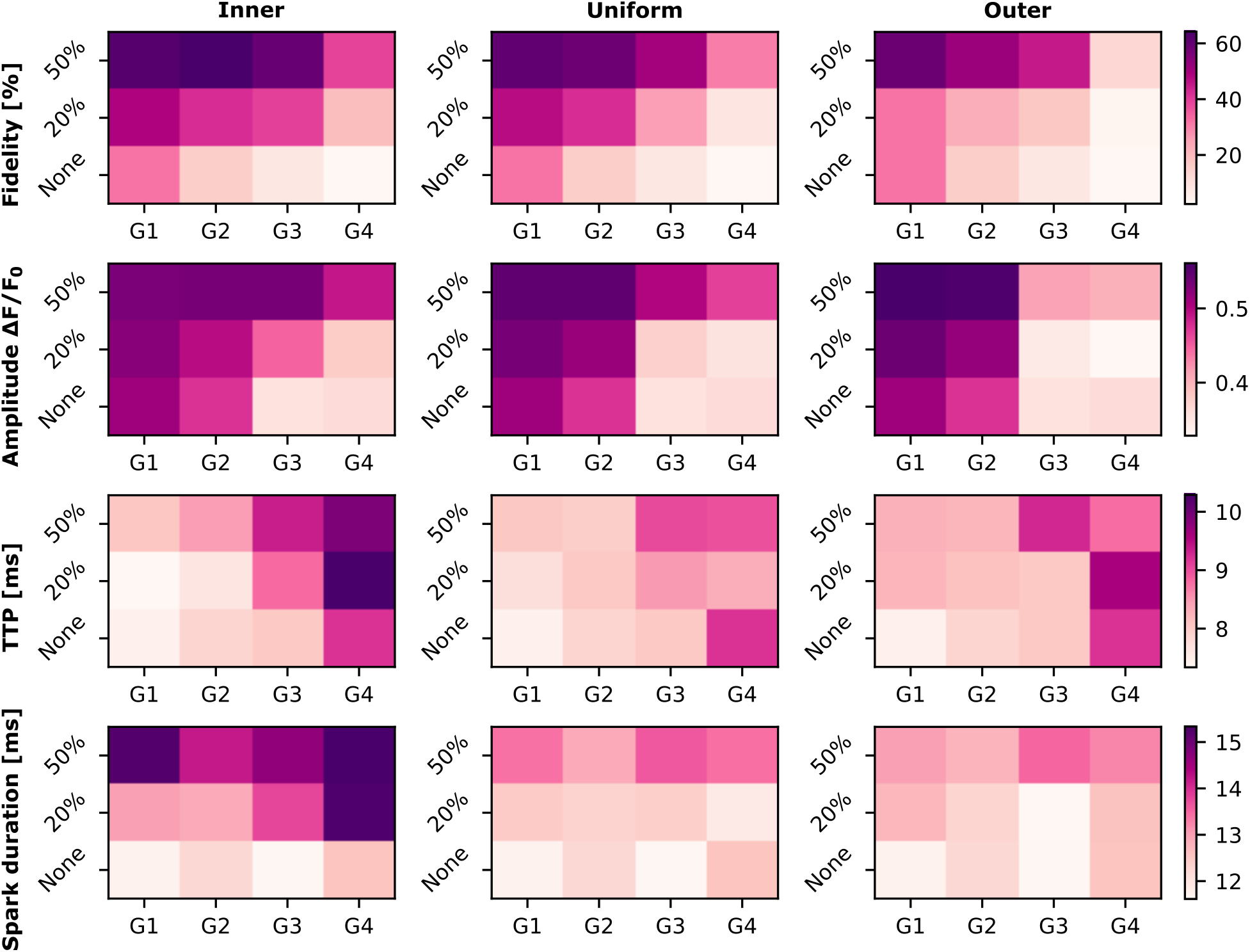
Heat maps depicting a summary of the Ca^2+^ spark properties for different phosphorylation patterns. The rows show the different spark properties (fidelity, amplitude, TTP and spark duration) that were analyzed. The columns represent the different phosphorylation patterns studied in this work. Ca^2+^ spark properties depend on RyR cluster geometry, cluster integrity, and spatial organization of phosphorylation.

An additional controversial aspect is the impact of PKA *versus* CaMKII on Ca^2+^ spark properties. Guo *et al.* (*2*) studied the effect of CaMKII on Ca^2+^ sparks in mouse ventricular myocytes. They observe an increase in Ca^2+^ spark frequency, spark duration, spatial spread, and amplitude due to CaMKII-mediated RyR phosphorylation. Furthermore, Li *et al.* (*29*) found that PKA increases Ca^2+^ spark frequency, amplitude, duration, and width in mouse ventricular myocytes. However, they conclude that PKA mediated changes in spark dynamics may be attributable to phospholamban and its resulting effects on SERCA rather than RyR phosphorylation (*29*). CaMKII and PKA therefore have different impacts on calcium sparks, although both phosphorylating proteins increase the open probability of the RyR (*29*). Further computational studies investigating phosphorylation patterns for CaMKII-specific sites may be of interest to determine whether distinct phosphorylation patterns occur in CaMKII-mediated phosphorylation and to understand why CaMKII and PKA show different phosphorylation effects as reported by Guo *et al.* (*2*) and Li *et al.*. For example, the larger size of CaMKII (56-58 kDa (*30*)) may hinder phosphorylation activity in the central part of the CRU in the narrow dyadic cleft, leading to an outer phosphorylation pattern.

In this study we reaffirm the importance of cluster geometry on Ca^2+^ spark properties. Additionally, these results show the large impact that spatial phosphorylation pattern can have on Ca^2+^ spark properties, leading to differences in spark fidelity, amplitude or duration depending on the pattern. Thus, it is necessary to conduct further imaging studies based on the spatial distribution of the phosphorylated RyRs as shown in (*8*) and on the relation between cluster geometry and phosphorylation during HF (*7, 28*). Differences in cluster geometry and spatial phosphorylation patterns could potentially explain the conflicting results from different studies on the impact of phosphorylation proteins on Ca^2+^ sparks (*24*), but also emphasize the need for future investigation of the role of PKA and CAMKII phosphorylation of RyRs during health and disease.

## Abbreviations

(ECC): excitation-contraction coupling
(SR): sarcoplasmic reticulum
(RyRs): ryanodine receptors
(SERCA): SR Ca^2+^-ATPase
(PKA): protein Kinase A
Ca^2+^ calmodulin kinase type II: (CaMKII)
(jSR): junctional SR
(nSR): non-junctional SR
(PCA): principal component analysis
(TTP): time to peak

## 5 Acknowledgements

MHM and KJM are supported by the Simula-UCSD-University of Oslo Research and PhD training (SUURPh) program, an international collaboration in computational biology and medicine funded by the Norwegian Ministry of Education and Research. P.R. would like to acknowledge support of this work by the Wu Tsai Human Performance Alliance and the Joe and Clara Tsai Foundation. This work used computational resources on the Oakforest-PACS supercomputer system provided by The University of Tokyo through Joint Usage/Research Center for Interdisciplinary Large-scale Information Infrastructures and High Performance Computing Infrastructure in Japan (Project IDs: JHPCN-jh180024).

## 6 Supplemental material

**Table 1:**
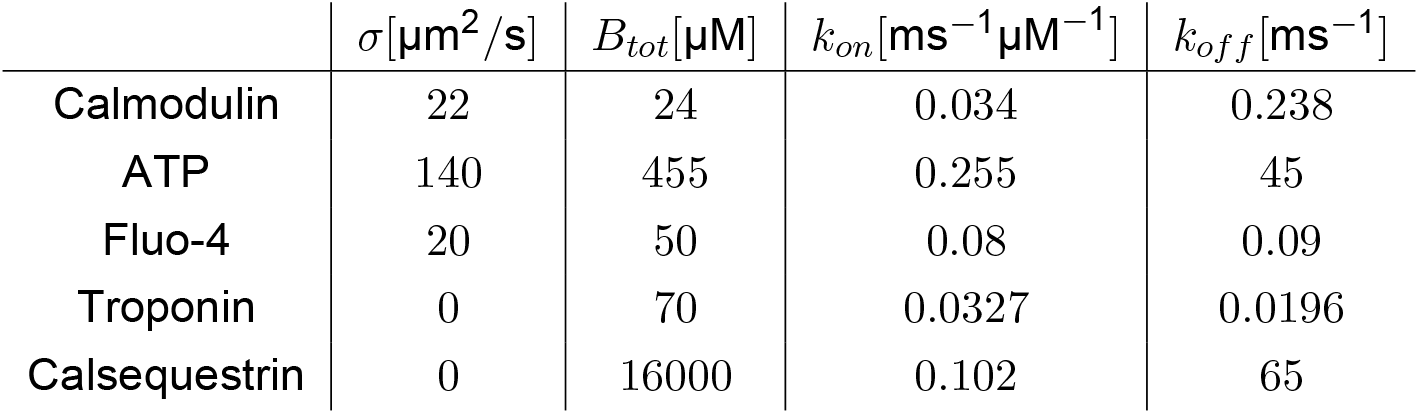
Buffer parameters.

**Table 2:** RyR and pRyR Model parameters.

| | $k_{min}[\text{ms}^{-1}]$ | $k_{max}[\text{ms}^{-1}]$ | $K_{RyR}^+[\mu\text{M}]$ | $K_{pRyR}^+[\mu\text{M}]$ | $n$ |
| --- | --- | --- | --- | --- | --- |
| + | $5 \times 10^{-6}$ | 0.9 | 55 | 25 | 2.7 |
| - | 0.9 | 3 | 35 | 35 | -0.5 |

**Table 3:** SR ratio and SERCA surface for the different geometries.

|  | G1 | G2 | G3 | G4 | G5 |
| --- | --- | --- | --- | --- | --- |
| jSR/nSR ratio | 26.19% | 27.53% | 29.67% | 32.37% | 40.48% |
| SERCA surface | $4.54 \mu\text{m}^2$ | $4.55 \mu\text{m}^2$ | $4.57 \mu\text{m}^2$ | $4.62 \mu\text{m}^2$ | $4.84 \mu\text{m}^2$ |

**Movie 1:** Simulation movie for G2, uniform phosphorylation.

**Movie 2:** Simulation movie for G2, inner phosphorylation.

**Movie 3:** Simulation movie for G2, outer phosphorylation.

**Movie 4:** Simulation movie for G3, uniform phosphorylation.

**Movie 5:** Simulation movie for G3, inner phosphorylation.

**Movie 6:** Simulation movie for G3, outer phosphorylation.

**Figure S1:**
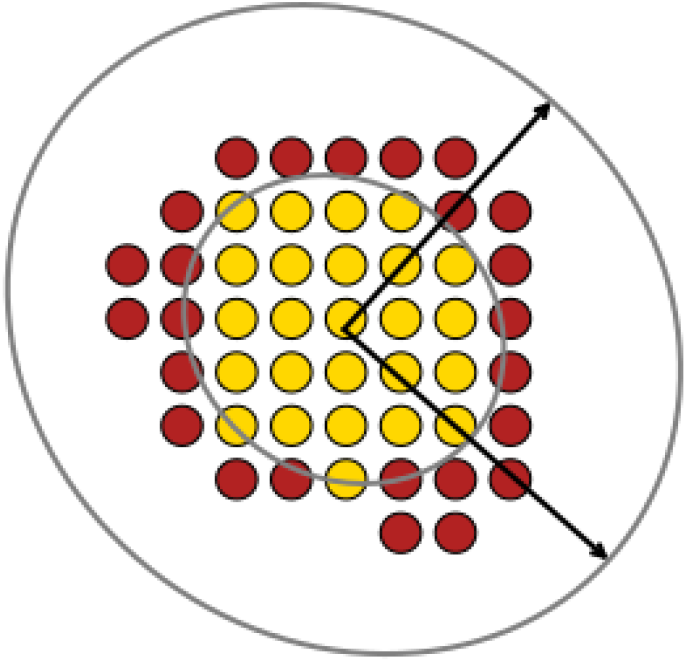
Schematic of the PCA algorithm for estimating the inner phosphorylation pattern. The arrows are vectors showing the eigenvectors of the covariance matrix of the spatial distribution of the RyRs. A first ellipse around the eigenvectors is shown. The ellipse dimensions are decreased until the desired number of phosphorylated RyRs is reached (in this case 50%).

**Figure S2:**
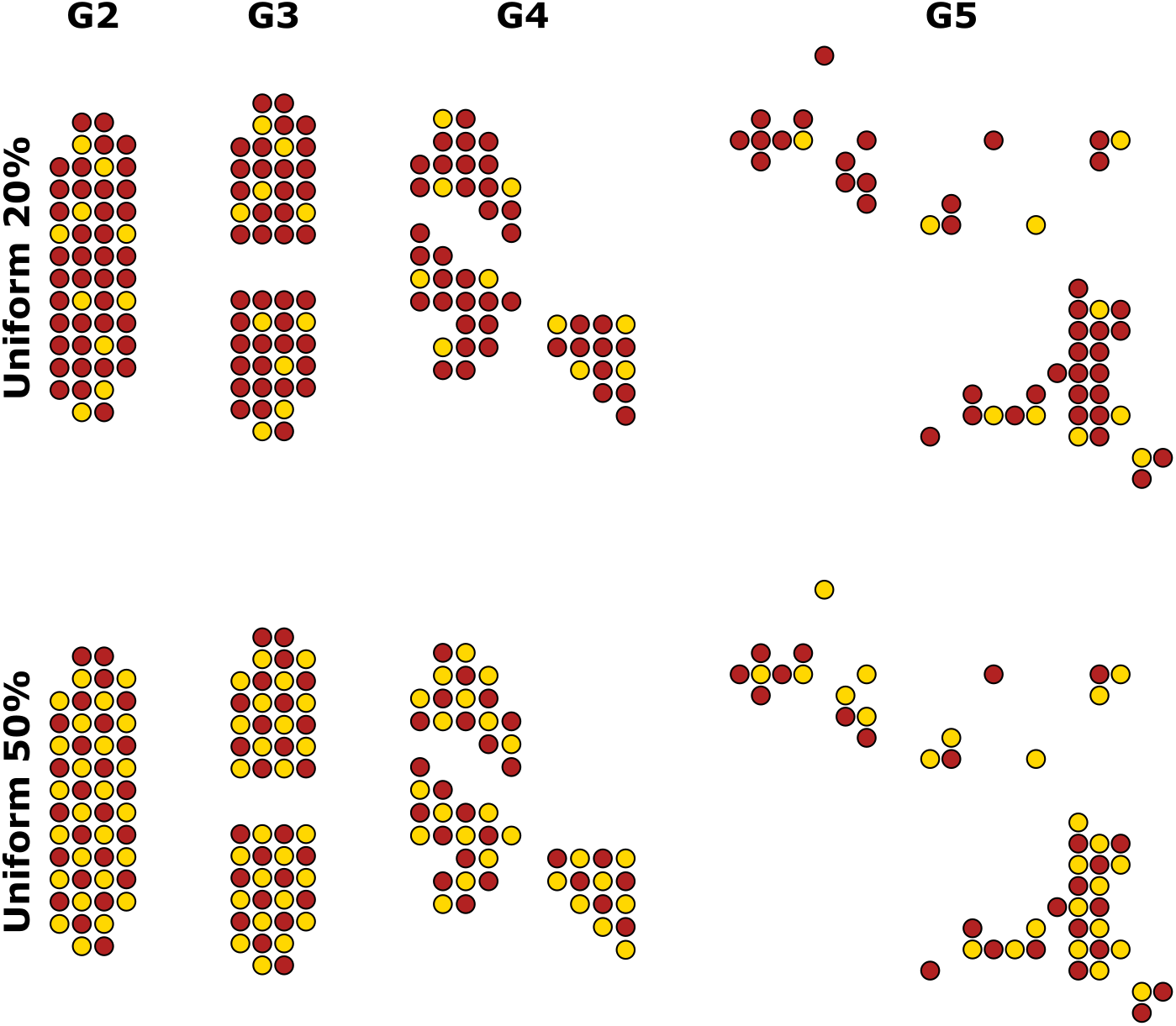
Uniform phosphorylation patterns for geometries G2-G5.

**Figure S3:**
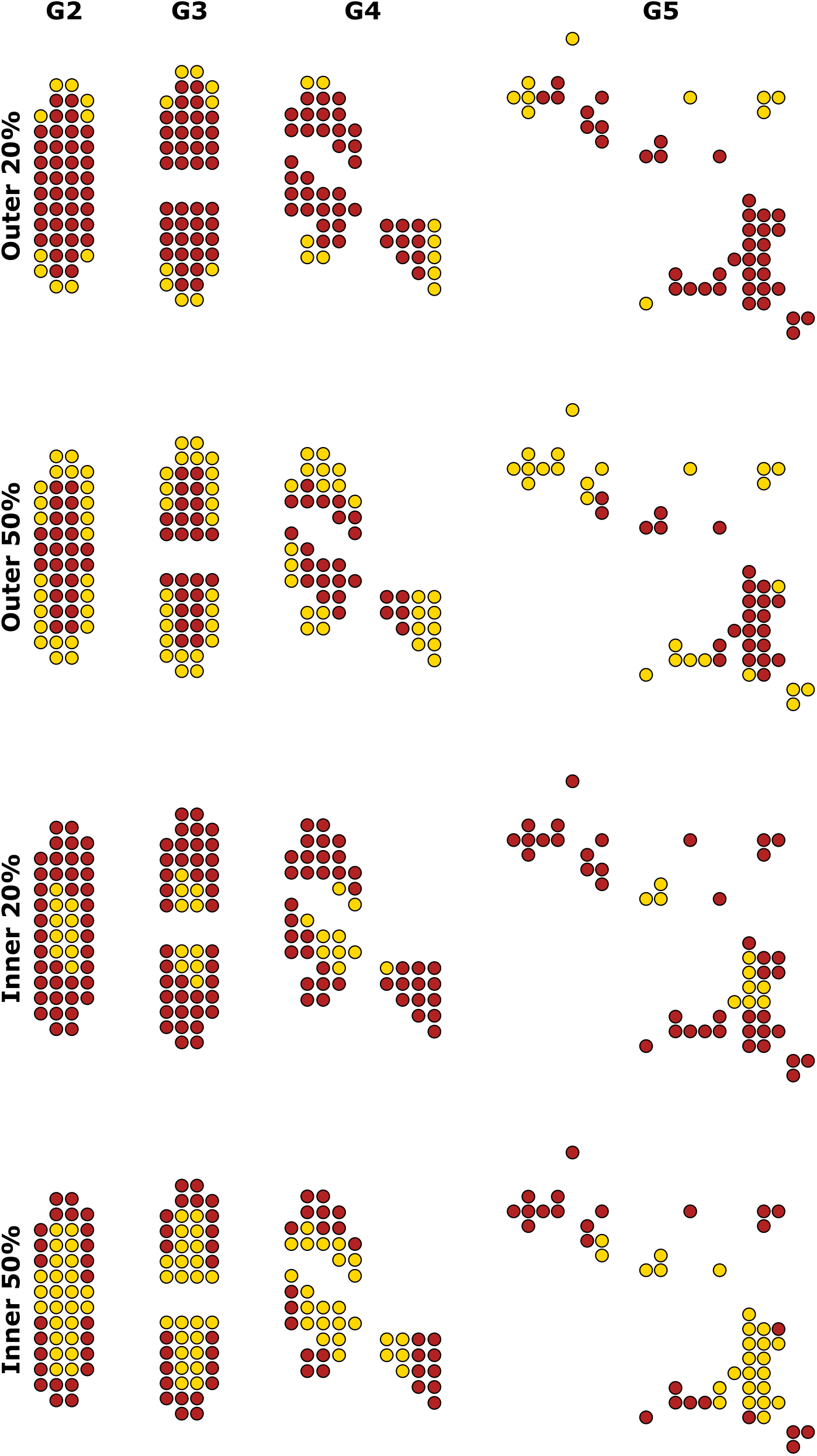
Inner and outer phosphorylation patterns for geometries G2-G5.

**Figure S4:**
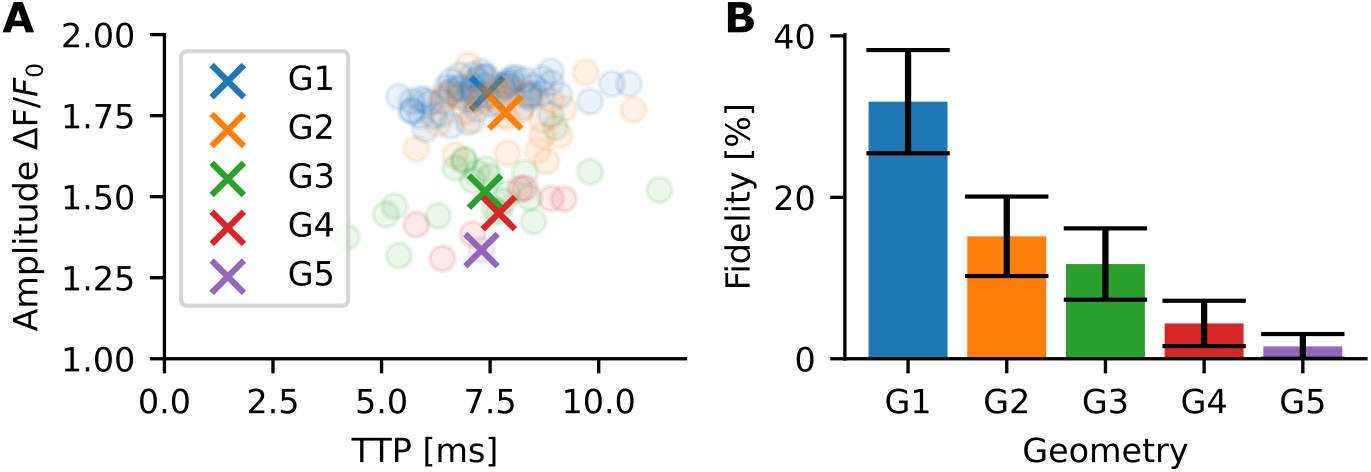
Spark properties of unphosphorylated RyR for different geometries. For each geometry 200 simulations were conducted. (A) The distribution created by the relation between TTP and amplitude is shown in a scatter plot shows. The crosses represent the mean values across all simulations. (B) The spark fidelity for each geometry is presented in a bar chart.

**Figure S5:**
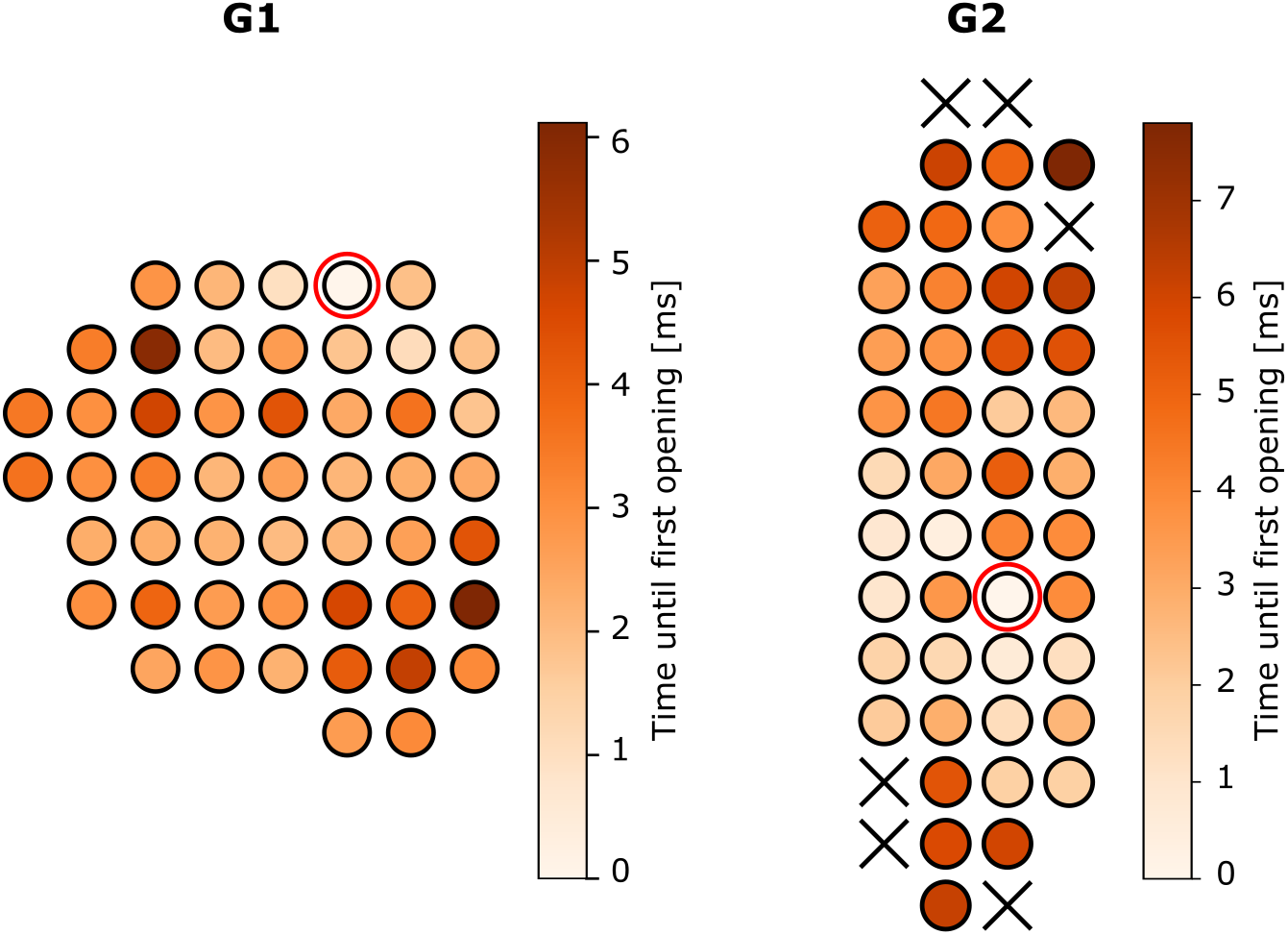
Activation map of two single simulations for G1 and G2 in the unphosphorylated state. The red circle indicates which RyR was opened to start the simulation.

